# Recombination Rates Are Governed by Sex-Specific Evolutionary Programs in House Mice

**DOI:** 10.1101/2025.06.16.659996

**Authors:** Lydia K. Wooldridge, Micah Pietraho, Peyton DiSiena, Sam Littman, Benjamin Clauss, Beth L. Dumont

## Abstract

Recombination rates vary markedly across species, populations, and sexes. In house mice (*Mus musculus*), this variation is particularly pronounced. Prior studies have established large differences in global recombination rates between *M. musculus* subspecies and inbred strains, with males exhibiting more extensive variation than females. The observation of sex-limited variation has prompted the hypothesis that male and female recombination rates may evolve by distinct evolutionary mechanisms in *M. musculus*. Here, we set out to formally evaluate this hypothesis in a phylogenetic framework using a dataset of cytogenetic sex-specific genome-scale crossover rate estimates from >6000 single meiotic cells from 31 genetically diverse inbred mouse strains spanning five *Mus* species and four *M. musculus* subspecies. Using phylogenetic comparative methods, we document a significant phylogenetic signal in male recombination rates, but female recombination rates show no clear phylogenetic trend. Males from *M. m. musculus* exhibit a large increase in recombination rate compared to other *M. musculus* subspecies, prompting us to explicitly test models of lineage-specific trait evolution. We show that the phylogenetic distribution of male recombination rates is best explained by an evolutionary model that allows a unique adaptive optimum along the *M. m. musculus* lineage, whereas female recombination rates are well-explained by a simplified model with a single global trait optimum. Taken together, our findings confirm the hypothesis that recombination rate evolution in house mice is governed by distinct sex-specific evolutionary regimes and motivate future efforts to ascertain the sex-specific selective pressures and sex-specific genetic architectures that underlie these observations.

**ARTICLE SUMMARY:** Meiotic recombination rates are highly variable between species, populations, and sexes. This variation is genetically controlled, but the underlying evolutionary processes that shape the extreme diversity of recombination rates are poorly understood. Here, we analyze sex-specific recombination rate estimates across a large panel of genetically diverse male and female house mice in an explicit phylogenetic framework. We show that recombination rates in males and females have evolved under distinct evolutionary programs, implying sex differences in the phenotypic value that optimizes evolutionary fitness. Our data point to intersexual genetic conflict driving rapid sex-specific recombination rate evolution in this system.

## INTRODUCTION

The rate of recombination is a fundamental parameter in genetics and evolutionary biology. It influences the evolutionary trajectories of new alleles within populations (Webster and Hurst 2012), governs the introduction of new haplotypes into the gene pool, and plays a central role in theories concerning the origin and maintenance of sexual reproduction (Otto and Lenormand 2002). Recombination also shapes various genomic characteristics, such as base composition (Duret and Arndt 2008) and levels of both DNA (Halldorsson et al. 2019; Palsson et al. 2025) and haplotype diversity (Pritchard and Przeworski 2001; Greenwood et al. 2004). It is also essential for ensuring the accuracy of meiosis: both excessively high and abnormally low recombination rates can lead to chromosome segregation errors, resulting in aneuploid gametes or early meiotic failure (Hassold and Hunt 2001).

A key principle of evolutionary biology is that functionally significant traits should be subject to stabilizing selection for a narrow range of optimal values. In perplexing contrast to this prediction, recombination rates display extensive variability across multiple evolutionary timescales (Coop and Przeworski 2007; Smukowski and Noor 2011; Peñalba and Wolf 2020). Across eukaryotes, global recombination rates span three orders of magnitude (Stapley et al. 2017). Even within mammals, genome-scale recombination rates range 10-fold (Dumont 2017a). Recombination rates also vary at the population level. For example, recombination rates in outbred mice (Dumont et al. 2009), humans (Cheng et al. 2009; Fledel-Alon et al. 2011), sheep (Johnston et al. 2016), and *Drosophila* (Hunter et al. 2016) differ by a factor of at least two.

In many species, recombination rate also exhibits striking sex dimorphism, or heterochiasmy. For example, in mammals, females typically have higher global recombination rates than males, but males exhibit localized enrichment of recombination in telomeres (Broman et al. 1998; Sardell and Kirkpatrick 2020). The causes of this sex dimorphism are largely unknown. The meiotic cells of the ovary and testis provide distinct, sex-specific milieus for recombination to unfold, establishing numerous plausible biological mechanisms for the emergence of sex differences in recombination. Indeed, sex differences in the temporal progression of meiosis (Hunt and Hassold 2002), meiotic chromatin structure (Gruhn et al. 2013), meiotic gene regulatory programs (Turner 2007), and meiotic checkpoint stringency (Morelli and Cohen 2005) could establish sex-specific evolutionary pressures on meiosis and recombination. Several theoretical hypotheses have also been advanced to explain heterochiasmy, including sex differences in the strength of epistasis (Lenormand 2003; Sardell and Kirkpatrick 2020), haploid selection (Lenormand 2003), meiotic drive (Brandvain and Coop 2012), and selection against gametic aneuploidy (Sardell and Kirkpatrick 2020). However, all presuppose that sex differences in recombination are adaptive (Sardell and Kirkpatrick 2020)—an assumption that has yet to be firmly established (Ritz et al. 2017)—and empirical support for these hypothesis remains limited.

We and others have previously established house mice (*M. musculus*) as an especially valuable model system for studying the causes of sex differences in recombination (Dumont et al. 2009; Liu et al. 2014; Peterson and Payseur 2021). Using both cytological and genetic approaches for quantifying genome-scale recombination rates, prior studies have shown that recombination rates are broadly conserved across female house mice, but markedly divergent across males of different house mouse strains and subspecies (Dumont and Payseur 2011; Peterson and Payseur 2021). Moreover, several inbred mouse strains have been identified that show a reversal in the most common direction of sexual dimorphism for recombination rate (Dumont and Payseur 2011; Peterson and Payseur 2021). For instance, males from several inbred strains of *M. m. musculus* have higher recombination rates than *M. m. musculus* females, contrasting the dominant trend of higher female recombination rates in strains from *M. m. domesticus* and *M. m. castaneus*. These directional shifts in heterochiasmy and the magnitude of male recombination rate divergence in house mice are surprisingly stark given that *M. musculus* subspecies diverged <500 thousand years ago (Phifer-Rixey et al. 2020).

While the cause(s) of these subspecies differences in recombination rate dimorphism is unknown, these observations would seem to suggest that male and female house mice evolve under different selective pressures or evolutionary regimes (Dumont 2011). Here, we set out to directly test this hypothesis using phylogenetic modeling. We compiled a set of cytogenetic recombination rate estimates for over 31 inbred mouse strains that capture each of the core *M. musculus* subspecies and 4 outgroup taxa. We then use whole genome sequences to reconstruct phylogenetic trees reflecting strain relationships and model distinct evolutionary scenarios in a phylogenetic framework. Our findings reveal distinct evolutionary processes shaping male and female recombination rates and suggest that adaptive mechanisms have contributed to patterns of male recombination rate divergence in *M. musculus*.

## METHODS

### Animal husbandry and experimental crosses

Wild-derived inbred strains CAROLI/EiJ, CZECHII/EiJ, GAIB/NachJ, JF1/MsJ, LEWES/EiJ, MANB/NachJ, MOLF/EiJ, PAHARI/EiJ, PERC/EiJ, PWK/PhJ, SARA/NachJ, SARB/NachJ, and SPRET/EiJ and were obtained from The Jackson Laboratory (JAX) Repository and housed in a low barrier room within the JAX Research Animal Facility. Strain Gor/TUA (RBRC01242) was cryorecovered from frozen embryos purchased from the RIKEN BioResources Center and maintained by sib × sib mating. Mice from inbred strain POHN/DehMmJax were originally obtained from Dr. David Harrison’s Laboratory at JAX and transferred into the Dumont Lab’s private colony. All mice were provided with access to food and water ad libitum. Sexually mature males were euthanized by exposure to CO_2_ at 8-26 weeks of age. Pregnant females were euthanized by CO_2_ inhalation at 16.5-18.5 days post coitus. Pups were then dissected out of the pregnant female uterus and sacrificed by decapitation with sharp blades.

All mice were housed and handled in strict accordance with an animal care and use protocol approved by the JAX Animal Care and Use Committee (Protocol # 17021).

### Assaying autosomal crossover rate via cytogenetic analysis of MLH1 foci in pachytene cells

Meiotic cell spreads were prepared from adult testis tissue and fetal ovarian tissue as previously described (Peters et al. 1997) and immunostained following published protocols (Dumont and Payseur 2011). Slides were blocked in blocking media [1% donkey serum, 3% bovine serum albumin (300mg/100mL; Fraction V; Fisher Scientific), and 0.0005% Triton X-100 v/v in 1× PBS (pH = 7.4)] and antibodies were diluted in antibody dilution buffer (ADB; 1% normal donkey serum, 0.3% bovine serum albumin v/v in 1× PBS). The following primary antibodies were used at the specified dilutions: 1:75 mouse anti-MLH1 (BD cat# 550838), 1:300 rabbit anti-SCP3 (Novus Biologicals cat # NB300-231), and 1:100 human anti-centromere polyclonal (Antibodies, Inc, cat # 15-234). The following secondary antibodies were used at 1:200 dilution: donkey anti-mouse Alexa Fluor 488, donkey anti-goat Alexa Fluor 594, and donkey anti-human Coumarin AMCA (Jackson Immunoresearch). Slides were mounted in ProLongGold antifade (Promega) and imaged at 63x on a Leica DM6B upright epifluorescent microscope equipped with DAPI, GFP, and Texas Red fluorescent filters and a cooled monochrome 2.8-megapixel digital camera. Images were post-processed and analyzed in the Fiji software package (Schindelin et al. 2012).

A minimum of 40 late pachytene cells characterized by (i) the complete merger of SYCP3 signals from all autosomal homologs; (ii) a full complement of chromosomes; (iii) low background fluorescence; and (iv) bright, punctuate MLH1 signals were imaged for each inbred strain. Cells that were damaged during cell spreading or displayed bulbous SYCP3 signals at chromosome termini (indicative of transition into diplotene) were excluded. For each cell meeting these criteria, the total number of autosomal MLH1 foci was recorded. MLH1 foci on the sex chromosome bivalent were excluded from all analyses, as the meiotic dynamics of the XY sex chromosomes are temporally decoupled from those of the autosomes (Kauppi et al. 2011; Acquaviva et al. 2020). In total, MLH1 counts were obtained for 1266 cells across 57 individuals representing 15 strains. These totals include 15 females from five strains (Gor/TUA, PERC/EiJ, POHN/DehMmJax, PWK/PhJ, and SARB/NachJ), and 42 males representing 14 strains (**Table S1**).

### Compilation of MLH1 count data

We combined our newly collected MLH1 counts with previously published MLH1 count data for genetically diverse inbred and outbred male and female mice (Lynn et al. 2002; Dumont and Payseur 2011; Peterson and Payseur 2021) (**Table 1; Table S1**). We limit our focus to wild-derived inbred strains, as the origin history of classical inbred mouse strains is complex and laboratory strains are unnatural hybrids carrying genomic ancestry from each of the three cardinal house mouse subspecies (*M. m. domesticus*, *M. m. castaneus*, and *M. m. musculus*) (Beck et al. 2000; Yang et al. 2007a). Cells with <19 or >44 MLH1 foci were excluded on the basis that a minimum of one crossover per autosome pair is required for accurate meiotic segregation (Page and Hawley 2003) and that cells with extreme numbers of MLH1 foci could represent staining artifacts. Within the Peterson and Payseur dataset, we further excluded cells with quality scores >4. Overall, the combined data set includes MLH1 counts from 6277 pachytene-stage cells from 31 strain backgrounds spanning five *Mus* species. The species *M. musculus* is represented by multiple inbred strains from each of the three primary *Mus musculus* subspecies (*M. m. domesticus*, *M. m. musculus*, *M. m. castaneus*), as well as multiple inbred strains of *M. m. molossinus*, a natural hybrid between *M. m. musculus* and *M. m. castaneus* (Yonekawa et al. 1988).

**Table 1.**
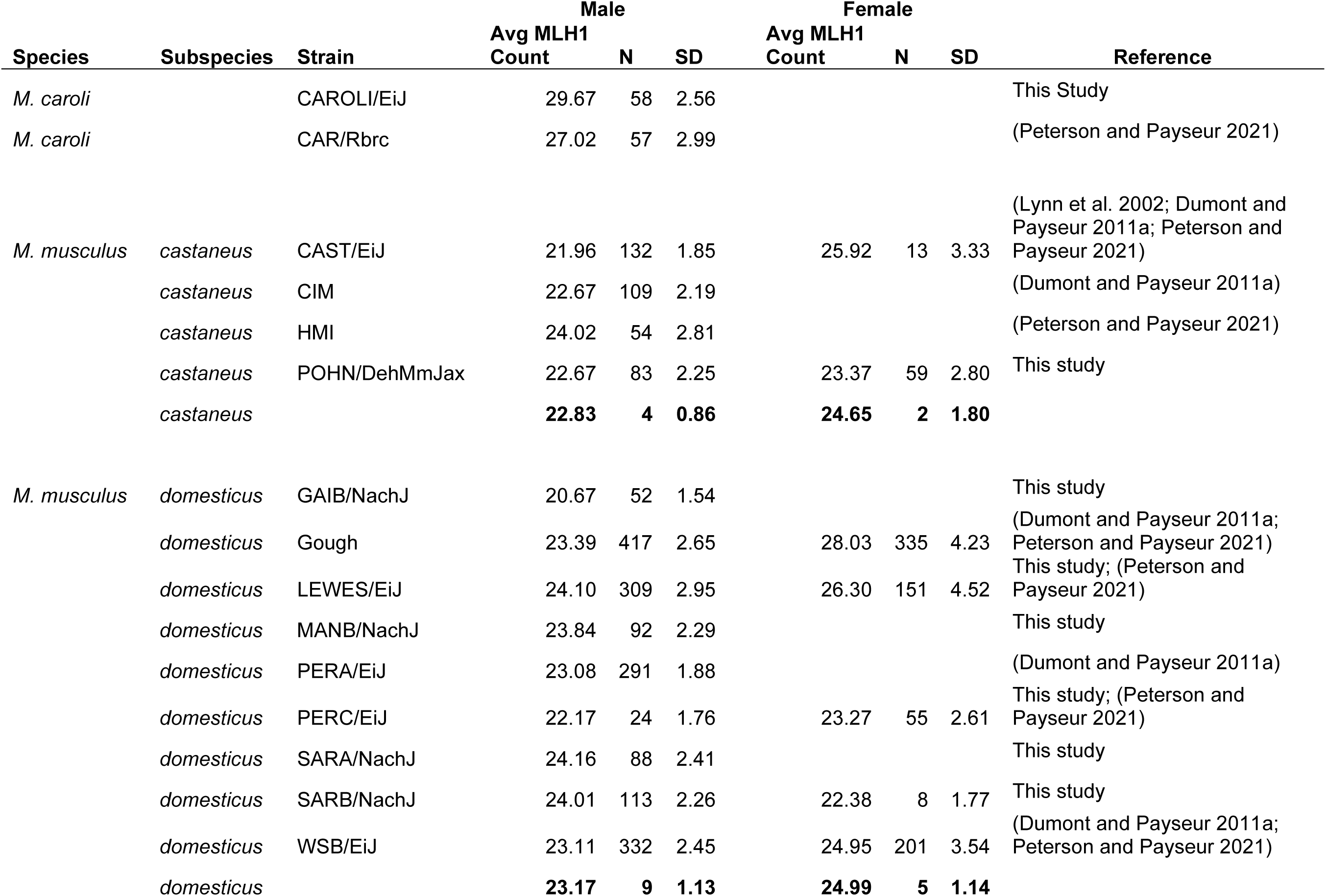
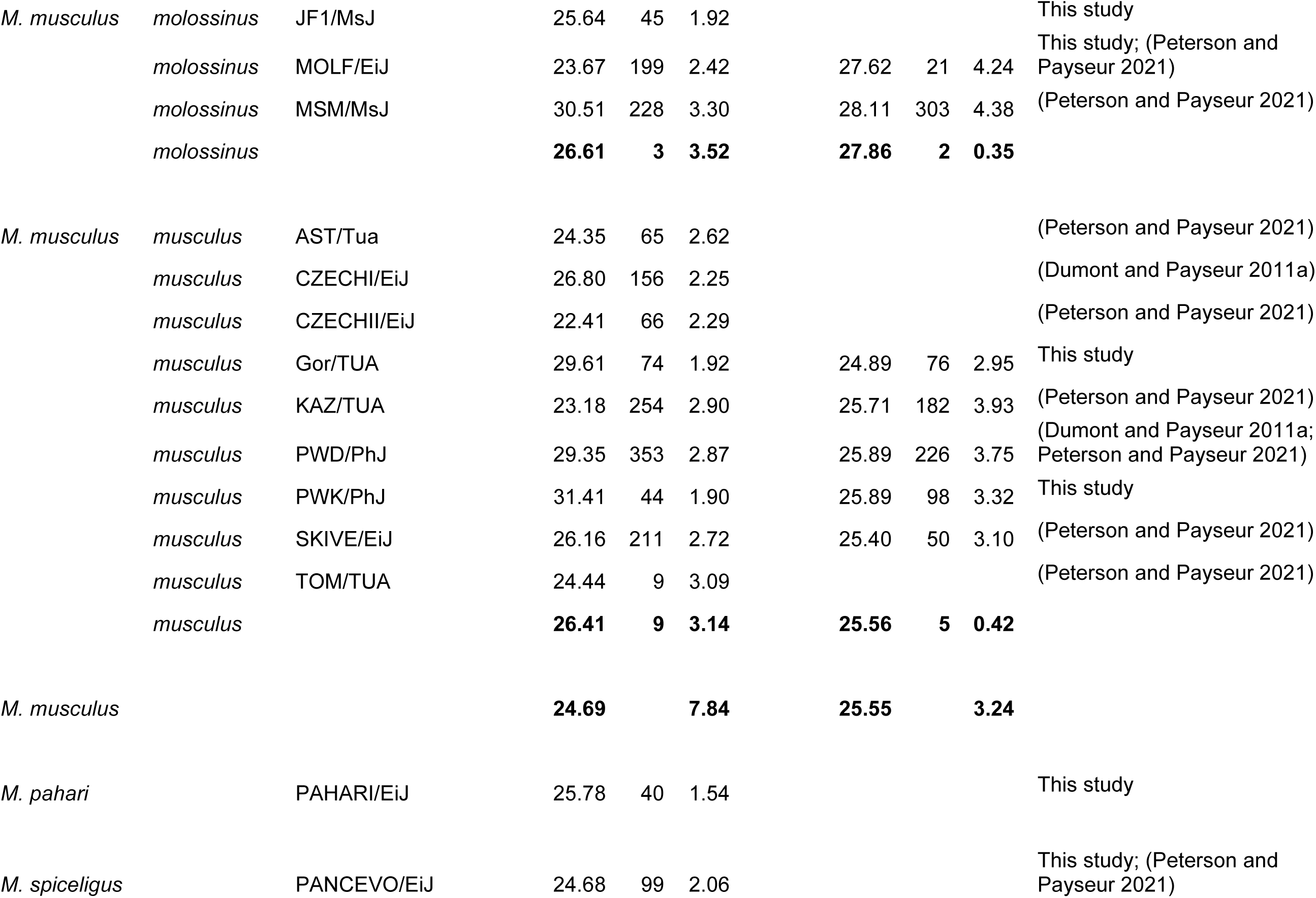
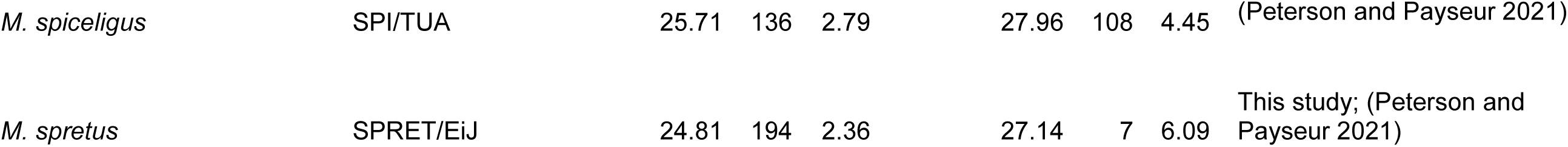
Sex-specific MLH1 foci counts in genetically diverse *Mus* strains.

Data for several strains were collected by different investigators in independent studies. In these circumstances, we confirmed consistency across scorers using an unpaired Wilcoxon-signed rank test. While some strains yielded significantly different MLH1 counts between studies, differences were numerically small in magnitude (<5.5%) and likely reflect study differences in treatment of MLH1 foci on the sex chromosomes or strain drift over the 10-year interval between the two studies (**Figure S1**, **Table S2**).

**Figure S1.**
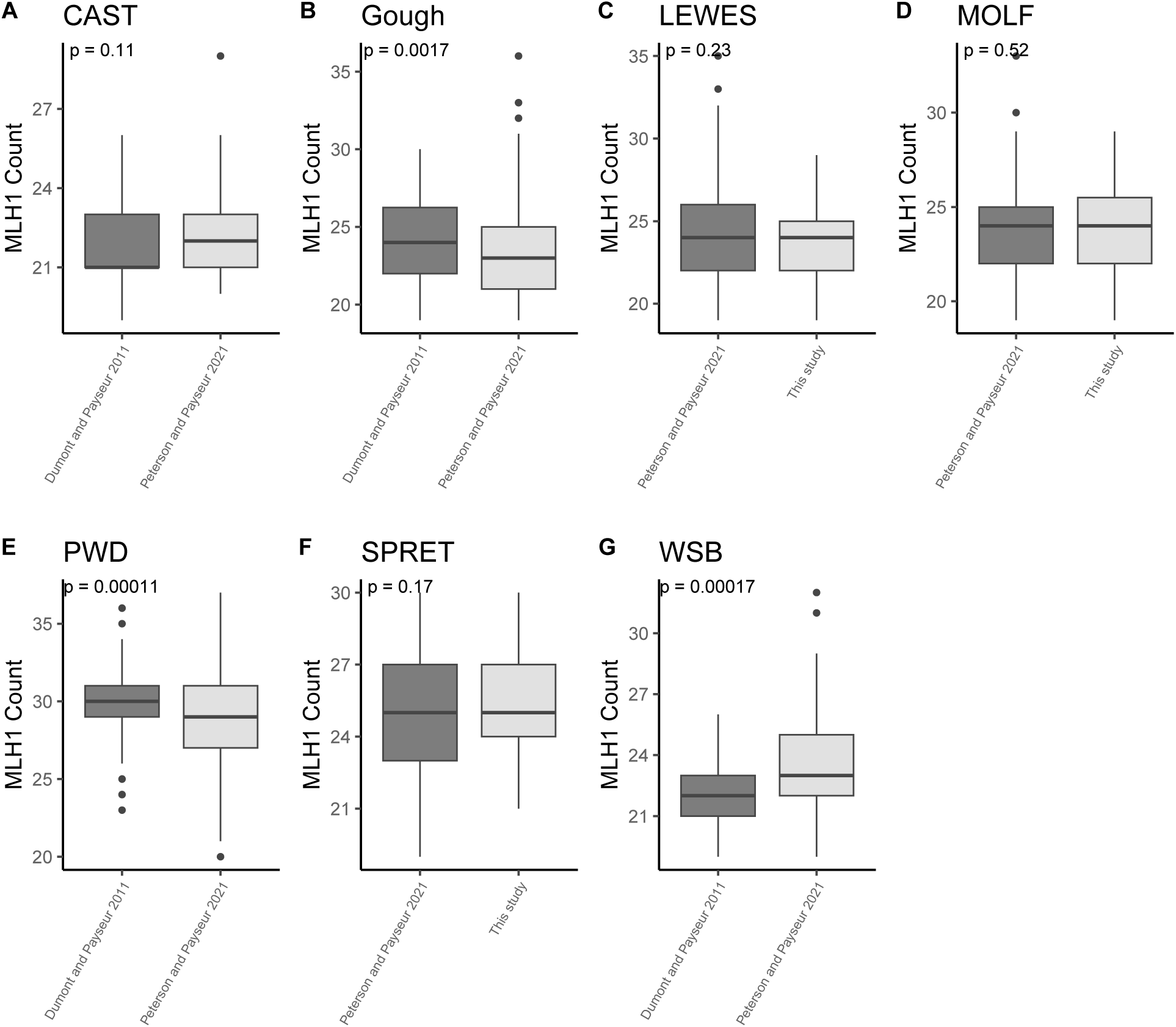
Comparison of MLH1 foci counts across independent studies. Boxplots display the number of autosomal MLH1 foci in spermatocytes from strains (A) CAST/EiJ, (B) outbred Gough island mice, (C) LEWES/EiJ, (D) MOLF/EiJ, (E) PWD/PhJ, (F) SPRET/EiJ, and (G) WSB/EiJ. *P*-values were calculated from Mann-Whitney U-tests. Significant, but quantitatively modest, differences in MLH1 foci counts are observed for Gough, PWD/PhJ, and WSB/EiJ. Note that Gough mice profiled in the Peterson and Payseur 2021 were significantly inbred, while those examined in Dumont and Payseur 2011 were outbred.

### Genome Sequencing and Genomic Data Analysis

Publicly available whole genome sequencing data are available for 18 of the 31 inbred strains included in our MLH1 dataset (**Table S3**). Three additional inbred strains (GAIB/NachJ, Gor/TUA, and POHN/DehMmJax) were *de novo* whole genome sequenced to ∼30x coverage using paired-end 150bp Illumina sequencing (Gor/TUA and POHN/DehMmJax) or ∼10x coverage using PacBio HiFi. For 8 of the remaining 10 strains with no corresponding whole genome sequence, we mined public sequencing archives for a genome sequence of a wild-caught mouse sampled in close geographic proximity to the inbred strain founders. Accession numbers for these wild-caught proxies are provided in **Table S3**. We were unable to identify suitable sequences for two outgroup strains (CAR/TUA and SPI/TUA); these strains were excluded from genomic and phylogenetic analyses described below.

The compiled genome sequences comprise a mixture of Illumina paired-end, BGI-SEQ, and PacBio HiFi sequencing reads. In all cases, fastq files were accessed from their respective sources and subjected to adaptor read trimming, file splitting, and quality control with fastp (v. 0.23.4). Reads were subsequently mapped to the GRCm39 reference mouse genome with bwa (v. 0.7.18; Illumina, BGI-SEQ) or pbmm2 (v. 2_1.16; HiFi). File merging and indexing were performed using samtools (v. 1.21), and duplicate reads were marked using samblaster (v 0.1.24). Bam files for samples sequenced across multiple libraries (PAHARI/EiJ and CAROLI/EiJ) were subsequently merged using samtools (v. 1.21). Per sample variant calling was performed using DeepVariant in WGS mode (v. 1.6.1; Illumina, BGI-SEQ) or HiFi mode (PacBio HiFi samples). Per sample gVCF files were then merged using glnexus (v1.4.1) under the DeepVariantWGS configuration to produce a joint call set. Sites were then filtered using bcftools (v. 1.16) to include only autosomal biallelic single nucleotide variants with ≤10% missing data and genotype quality ≥30. All bioinformatic analyses were performed using containerized software on the JAX High Performance *Sumner2* compute cluster.

### Phylogenetic Tree Construction

Phylogenetic trees were constructed from the filtered joint call set. Briefly, variants were greedily thinned to include only those sites with *r^2^* > 0.2 using PLINK (v2.00a2.3LM). The thinned VCF file was then converted into FASTA format using a custom perl script (vcf_to_fasta_justSNPs.pl). Subsequent conversions from FASTA to Stockholm and Phylip alignment formats were made using the SeqIO module for BioPython (version 3.12.9).

Quicktree (v. 2.0) was used to build a neighbor joining tree from the LD-thinned SNPs. We used the same SNP dataset to infer a maximum likelihood (ML) phylogenetic tree using phyml (v. 3.3). We specified a GTR model of nucleotide evolution, nucleotide frequencies computed from the empirical data, and estimated the transition/transversion ratio, proportion of invariant sites, and gamma distribution of rate classes via maximum likelihood. The executed command was:

phyml --input <ALIGNMENT FILE> \

--datatype nt \

--bootstrap -1 \

--model GTR \

-f e -t e \

--pinv e --alpha e \

--r_seed 12345

### Phylogenetic Modeling

Under a neutral (i.e., Brownian motion) model of evolution, the extent of phenotypic divergence between taxa should be proportional to their genetic divergence. In this circumstance, the underlying phylogenetic tree relating samples ought to be a good predictor of the distribution of trait values across taxa. To evaluate this hypothesis, we applied a mixed model approach to estimate the proportion of variation in sex-specific recombination rates that is attributable to the underlying sample phylogeny (Lynch 1991). Specifically, the model partitions each realized phenotypic value, *z*, into three components

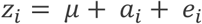

where 𝜇 is the phenotypic contribution shared by all members of the phylogeny, *a_i_* is the heritable additive effect of the trait in the *i*^th^ taxon, and *e_i_* is a residual error term that captures phenotype variation due to measurement error, phylogenetic uncertainty, phenotypic plasticity, rapid evolution along terminal branches, and fluctuating selection. Model fitting was performed via the compar.lynch function call with the ape package for R (Paradis et al. 2004; Paradis and Schliep 2019). ML parameter estimates were obtained using the EM algorithm, with convergence established when successive iterations yielded parameter estimates that differed by less than 1×10^−4^. The variances associated with the heritable additive and error terms, 𝜎_a_^2^ and 𝜎^2^, were then combined to calculate the phylogenetic heritability (𝐻^2^) of both male and female recombination rates:

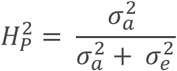

The significance of observed 𝐻^2^ values was assessed by an ad hoc permutation test. Recombination rates were randomly shuffled with respect to species designations at the tips of the tree, while the underlying topology and branch lengths remained constant. We then recomputed the phylogenetic heritability on the permuted dataset and determined the quantile position of our observed value along the distribution of 1000 permuted values.

To explicitly model lineage-specific adaptive shifts in recombination rate, we used the Ornstein-Uhlenbeck phylogenetic modeling framework implemented in the R package OUCH (Butler and King 2004; Cressler et al. 2015). OU models extend a neutral phylogenetic model of trait evolution (i.e., Brownian motion) through the inclusion of additional parameters that specify an optimal trait value and selection intensity along predefined lineages. For both male and female recombination rates, we compared a model with a unique trait optimum along the *M. m. musculus* lineage to null models of Brownian motion and a single phylogeny-wide trait optimum. Model selection was performed by comparison of corrected AIC values.

Code required to replicate these phylogenetic analyses is provided on FigShare (https://figshare.com/projects/Recombination_Rates_Have_Been_Shaped_by_Sex-Specific_Evolutionary_Programs_in_House_Mice/252902).

## RESULTS

### Species, Subspecies, Strain, and Sex diversity in global recombination rates

MLH1 is a mismatch repair protein that localizes to sites of crossing over. Although a subset of crossovers in mammals is resolved by a MLH1-independent pathway, this percentage is small (<10%) (Holloway et al. 2008; Gray and Cohen 2016). Thus, the localization of MLH1 on meiotic chromosome axes during prophase I reproduces the frequency and distribution of crossovers and can be used as a proxy for the genomic crossover distribution and rate (Anderson et al. 1999). We quantified global crossover rates via cytological evaluation of MLH1 foci in 1266 pachytene stage cells from 57 animals representing 15 genetically diverse inbred mouse strains. We combined these data with previously published MLH1 foci counts from house mice (see Methods) to assemble a large dataset of genomic crossover rate estimates from over 6,000 pachytene cells from 31 mouse strains representing five independent *Mus* species and four *Mus musculus* subspecies (**Table S1**; **Table 1**). This dataset includes MLH1 foci counts derived from both spermatoctyes (males) and oocytes (females) for 15 inbred mouse strains, providing a resource for directly testing hypotheses of sex-specific recombination rate evolution.

Species in our dataset span ∼5-7.5 million years of evolutionary divergence (Chevret et al. 2005; Thybert et al. 2018). Curiously, strains with the largest difference in MLH1 foci counts are not necessarily the most evolutionarily divergent. Across males, average MLH1 counts per strain range from 20.67 in the *M. m. domesticus* strain GAIB/NachJ to 31.41 in the *M. m. musculus* strain PWK/PhJ – two taxa that diverged <0.5 MYA (Phifer-Rixey et al. 2020). Considering only females, MLH1 counts range from 22.4 in SARA/NachJ to 28.1 in MSM/MsJ, strains that again derive from two closely related *M. musculus* subspecies. With the exception of *M. pahari*, which has a diploid chromosome number of 48, all *Mus* species in our dataset are represented by a conserved 2N=40 karyotype comprised of acrocentric chromosomes. Thus, the absence of a clear species-level effect on recombination rate divergence is not simply a consequence of evolutionary shifts in the chromosomal constraints on recombination (Dumont 2017a).

Prior work has established significant subspecies-level recombination rate divergence among males of the *M. musculus* species complex, with *M. m. musculus* males exhibiting markedly increased MLH1 counts compared to males from *M. m. domesticus* and *M. m.* castaneus (Dumont and Payseur 2011; Peterson and Payseur 2021). The addition of data from several new *M. musculus* strains lends further support for these trends (**Fig 1**; **Table 1**). On average, males from inbred *M. m. musculus* strains have higher MLH1 counts than males from either *M. m. domesticus* or *M. m. castaneus* (*musculus*: 26.41; *domesticus*: 23.17; *castaneus*: 22.83). Males from inbred strains of *M. m. molossinus* have similar MLH1 counts as *M. m. musculus* males (26.41 versus 26.61). *M. m. molossinus* is a natural hybrid between *M. m. musculus* and *M. m. castaneus* (Yonekawa et al. 1988), with genomic studies suggesting a dominant contribution of the *M. m. musculus* genetic background (Yang et al. 2007b). This suggests the possibility of shared genetic factors in *M. m. molossinus* and *M. m. musculus* contributing to increased male recombination rates in these subspecies.

**Figure 1.**
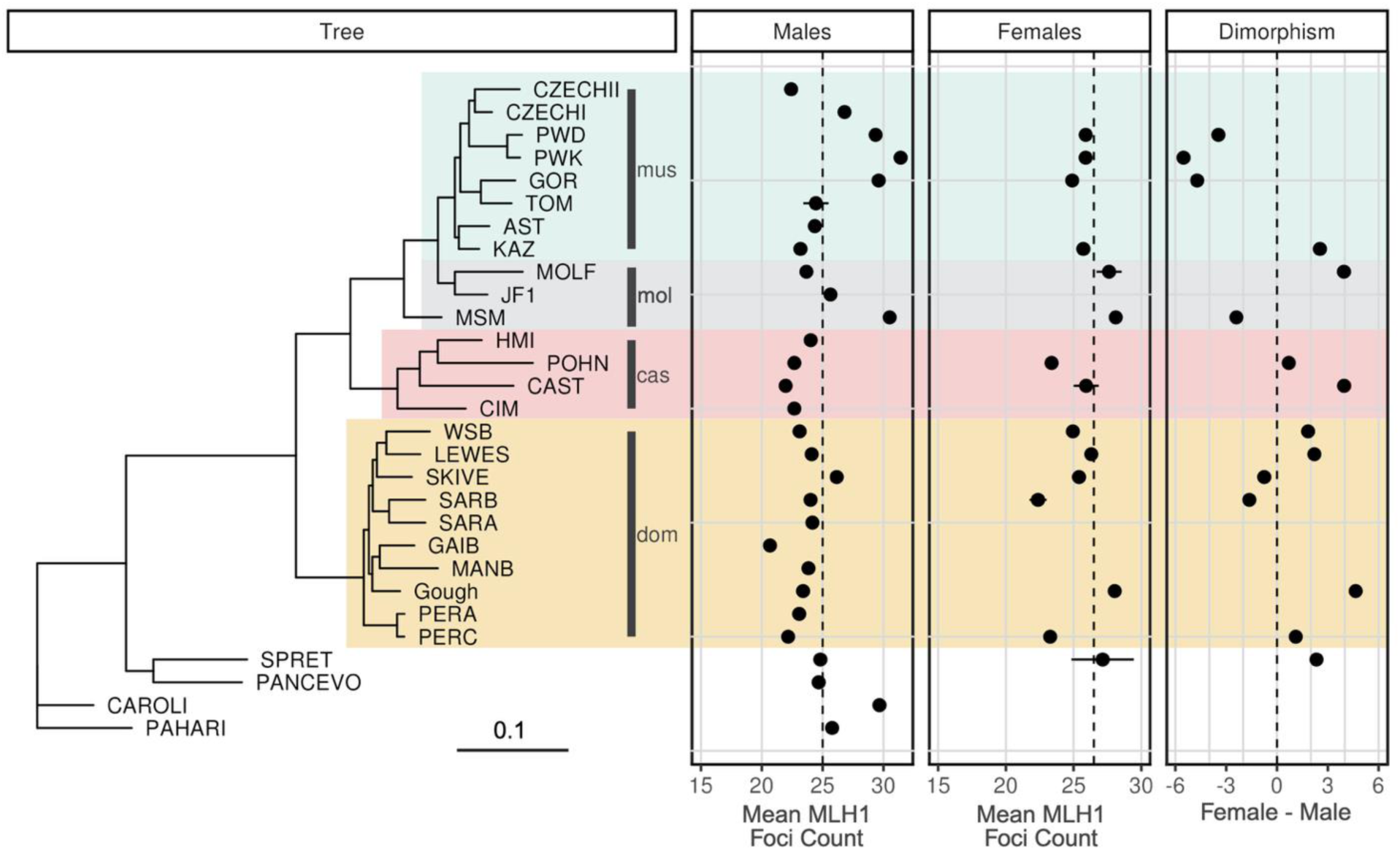
Maximum likelihood phylogenetic tree for genetically diverse inbred mouse strains. Tree is rooted to the outgroup species *M. pahari* represented by inbred strain PAHARI/EiJ. For brevity, strain labels at the tips of the tree exclude laboratory codes. Wild-derived inbred strains from *M. musculus* subspecies are labeled and color-coded in gold (*domesticus*), red (*castaneus*), gray (*molossinus*), and turquoise (*musculus*). Panels to the right of the tree plot average MLH1 foci counts (±1 standard error) for males and females for each strain, as well as the difference between male and female MLH1 foci counts. Note that MLH1 foci counts are not available for females from all strains. **Figure S2** presents a conceptually identical figure, but with a neighbor joining tree instead of a ML tree. Figure made using the ggtree package for R with aesthetic modifications in BioRender.

In contrast to the pronounced recombination rate divergence in *M. musculus* males, prior studies have revealed minimal divergence in recombination rate among females from the three cardinal house mouse subspecies (Dumont and Payseur 2011; Peterson and Payseur 2021). This conclusion is again reinforced by our addition of female MLH1 foci counts for several new inbred strains. The variance in average strain-level MLH1 counts for *M. musculus* females is smaller than the corresponding measure for males (3.24 versus 7.84), suggesting increased evolutionary constraint on female versus male recombination rates (**Table 1**).

Overall, average female MLH1 counts are higher than the corresponding values for males (26.5 versus 25.0; Mann-Whitney U-test, *P* = 3.57 x10^-38^), recapitulating the well-established pattern of heterochiasmy in house mice (Mallyon 1951; Dietrich et al. 1996; Shifman et al. 2006). However, at the level of individual inbred strains, there is considerable heterogeneity in the direction of this sex dimorphism. For example, male Gough mice have ∼4.6 fewer MLH1 foci than their female counterparts, whereas male PWK/PhJ mice have on average 5.5 MLH1 foci more than females (**Table 1**; **Figure 1**). Overall, the majority of *M. m. musculus* strains exhibit increased male MLH1 foci counts compared to females. In contrast, most *M. m. domesticus* and *M. m. castaneus* strains show the opposite pattern of increased female MLH1 foci counts. It is notable that the one *M. m. musculus* strain defying these predictions (i.e., increased female MLH1 foci counts compared to males) derives from Kazakhstan, a location close to the presumed ancestral region where the *M. musculus* species first emerged (KAZ/TUA; (Boursot et al. 1993a; Suzuki et al. 2013)). This observation tentatively suggests that the directional change in heterochiasmy in *M. m. musculus* occurred after initial subspecies divergence and migration out of the ancestral cradle.

**Figure S2.**
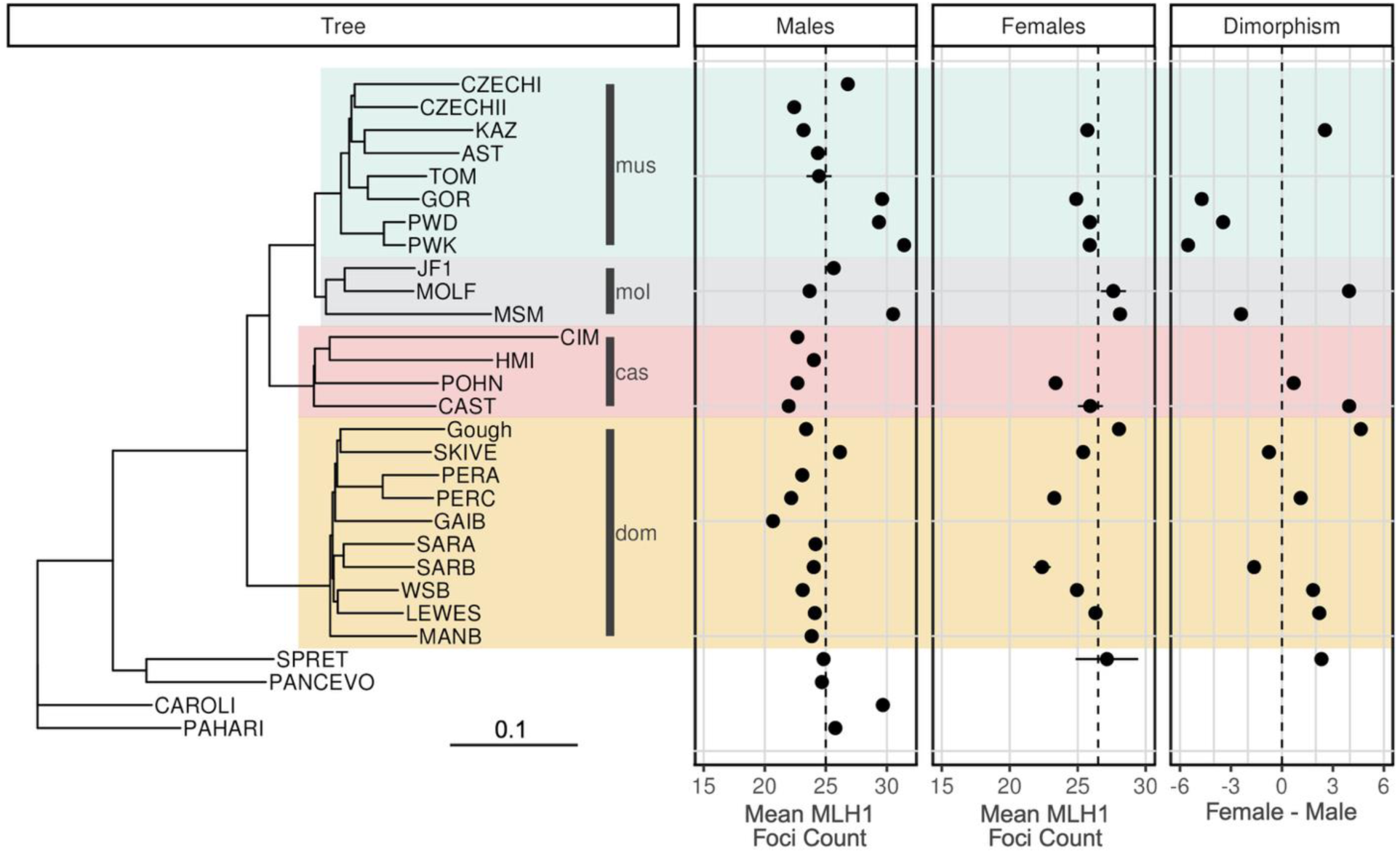
Neighbor joining phylogenetic tree for genetically diverse inbred mouse strains. Tree is rooted to the outgroup species *M. pahari* represented by inbred strain PAHARI/EiJ. For brevity, strain labels at the tips of the tree exclude laboratory codes. Wild-derived inbred strains from *M. musculus* subspecies are labeled and color-coded in gold (*domesticus*), red (*castaneus*), gray (*molossinus*), and turquoise (*musculus*). Panels to the right of the tree plot average MLH1 foci counts (±1 standard error) for males and females for each strain, as well as the difference between male and female MLH1 foci counts. Note that MLH1 foci counts are not available for females from all strains. Figure made using the ggtree package for R with aesthetic modifications in BioRender.

### Shared phylogenetic ancestry differentially accounts for variation in male and female MLH1 foci counts

We next set out to assess the distributions of MLH1 foci counts in male and female house mice in an explicit evolutionary framework using phylogenetic comparative methods and phylogenetic modelling. Under a model of neutral phenotypic evolution, the distribution of MLH1 foci counts across species should be well predicted by the underlying phylogeny relating strains. In contrast, if MLH1 foci counts are not genetically determined, are subject to strong stabilizing selection, or evolve exceptionally rapidly, strain variation in this phenotype may not mirror shared ancestry. To assess these possibilities, we first quantified the phylogenetic heritability (𝐻^2^) of MLH1 counts (Lynch 1991). The phylogenetic heritability is analogous to traditional broad sense heritability estimate: it represents the proportion of variation in a trait of interest that is explained by the underlying phylogeny relating the surveyed samples. Male MLH1 foci counts exhibit modest, but significant, phylogenetic heritability across a maximum likelihood (ML) tree of *Mus* species (𝐻^2^ = 0.46, *P* = 0.044). In contrast, 𝐻^2^ for female MLH1 foci counts is essentially zero, indicating that shared evolutionary descent does not predict the strain distribution of this trait (𝐻^2^ = 0.009, *P* = 0.614).

Recognizing that these discordant findings could reflect the increased number of strains with MLH1 foci count data for males compared to females, we pruned the ML tree and pared down our MLH1 dataset to include only the 15 strains with both male and female MLH1 foci counts (**Table 1**). We then re-estimated 𝐻^2^ on the thinned datasets. Our findings recapitulate those from the original, larger data (males, ML tree: 𝐻^2^= 0.45, *P* = 0.076; females ML tree: 𝐻^2^= 0.0097, *P* = 0.596). We next assessed the robustness of our results to implicit assumptions of the phylogenetic mixed model fit to estimate variance components. First, 𝐻^2^ estimation assumes that MLH1 counts are normally distributed. While the appropriateness of the normality assumption can be difficult to evaluate for small datasets, we find no overt departure from this assumption in our data (**Figure S3**). Second, the model assumes that phenotypes are estimated without systematic error. Mean MLH1 foci counts were computed across a minimum of 7 cells per strain, in most cases averaging over pachytene cells from multiple animals per strain to account for individual-level biological variation (**Table S1**). Further, given that the immunofluorescent assay for detecting MLH1 counts is identical for both spermatocyte and oocyte cell spreads, it is difficult to envision systematic sex biases in the technical error profile of the data used in our analysis. Third, phylogenetic heritability estimates assume that the tree specified in the model is correct. We performed phylogenetic inference using whole genome sequence data from analyzed strains or their close geographic proxies from the wild. The reliance on genomic data effectively integrates over regional fluctuations in evolutionary history due to incomplete lineage sorting or introgression, assuring that inferred trees are an accurate portrayal of species and strain relationships. Our tree also recapitulates known relationships among *Mus* species and clusters strains by subspecies and geographic origin (**Figure 1**), as expected (Chevret et al. 2005). Further, our findings are robust to the method of phylogenetic inference, as we recover qualitatively similar 𝐻^2^ estimates when accounting for ancestry using an NJ tree (male 𝐻^2^ = 0.62, P = 0.038; female 𝐻^2^ = 0.0097, P = 0.560). Lastly, and most obviously, the trees informing our 𝐻^2^ estimates for male and female MLH1 count data are identical. Thus, quantitative sex differences in 𝐻^2^ estimates cannot be explained by differences in the phylogeny.

Overall, our findings indicate that phylogenetic relatedness is a stronger predictor of male recombination rates than female recombination rates. This finding could be attributed to one or more explanations. First, male recombination rates may be determined by more stringent genetic controls than female recombination rates (*i.e.,* have higher genetic heritability). Second, female recombination rates may be more plastic or malleable than male recombination rates in response to environmental perturbations. Third, our findings could imply that female recombination rates are subject to more intense stabilizing selection across the *Mus* taxa evaluated here, with minimal additive trait variance in MLH1 counts leading to near-zero estimates of 𝐻^2^ Finally, the absence of a phylogenetic signal in females could be due to extremely rapid recombination rate evolution in this sex, with trait values fluctuating so rapidly in time that they override signals of shared evolutionary history. We discount this final interpretation as unlikely given the lower variance in MLH1 foci counts in females compared to males (**Table 1**).

**Figure S3.**
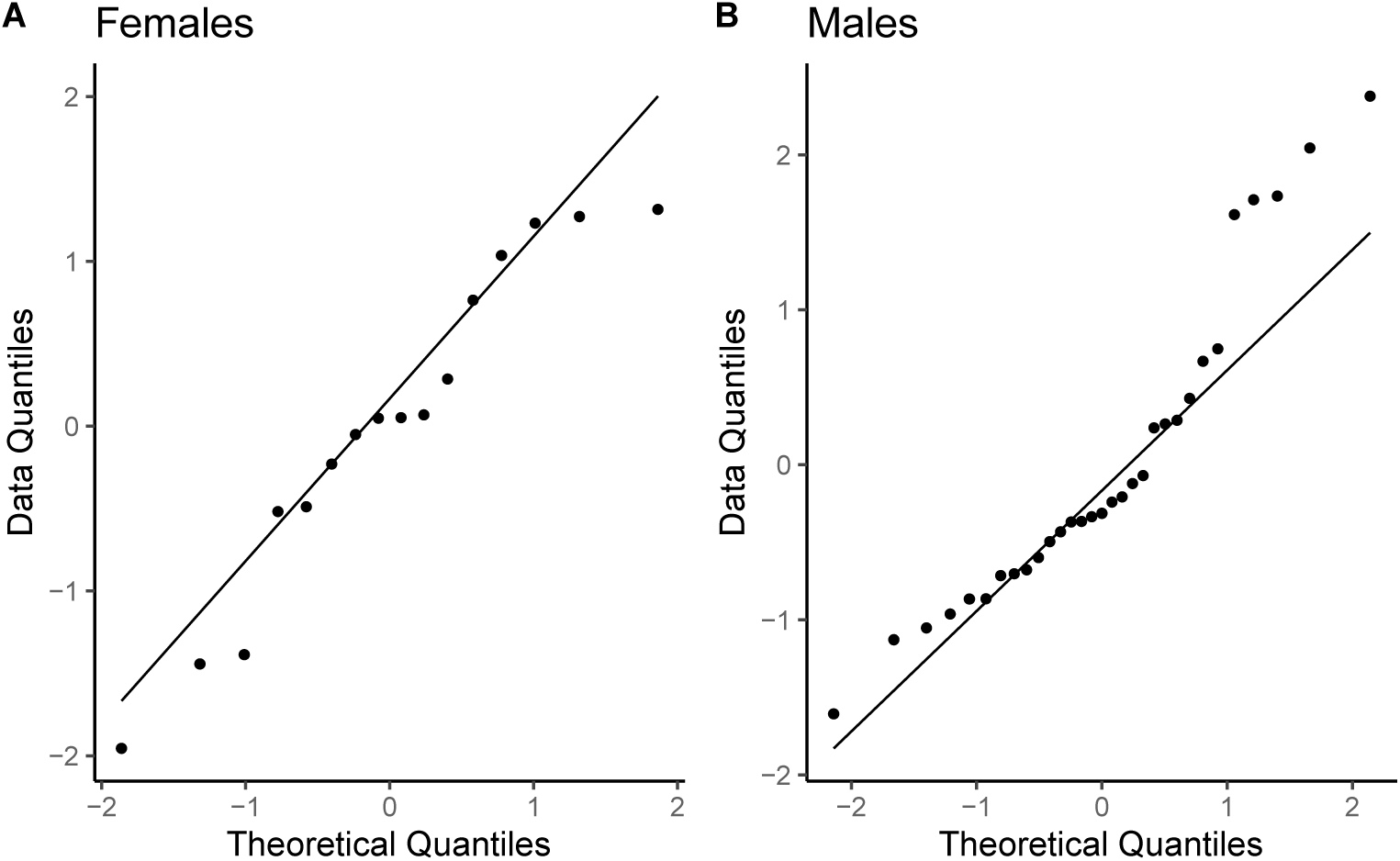
Quantile-quantile plots examining the relationship between theoretical quantiles from a normal distribution and observed sample distributions of (A) female and (B) male MLH1 foci counts. Data points largely fall along the y=x line, with some skewing at the extremes. Plots were generated in RStudio (v. 2023.09.1) using the geom_qq plotting function in ggplot.

### Modeling of sex-specific selection regimes for recombination rates in Mus musculus

Having discovered distinct phylogenetic trends in recombination rate evolution between males and females, we next sought to explicitly model defined scenarios of sex-specific recombination rate evolution in a phylogenetic context. We were particularly interested in evaluating the possibility of an adaptive increase in male recombination rates along the *M. m. musculus* subspecies lineage, given that males from this taxon consistently exhibit among the highest MLH1 focus counts and capture a shift in the direction of sex dimorphism for recombination rate (**Fig 1**). We modeled this evolutionary scenario as an Ornstein-Uhlenbeck process, specifying a unique adaptive trait optimum and evolutionary rate along the *M. m. musculus* lineage. We then compared this evolutionary model to null models wherein (1) the evolution of MLH1 values across all strains is determined by Brownian motion or (2) the evolution of MLH1 values is governed by selection toward a single, shared trait optimum.

A model with a unique adaptive optimum in *M. m. musculus* is a better fit to the phylogenetic distribution of male MLH1 focus counts than a Brownian motion model or model with a single global trait optimum, as assessed by the corrected Aikake Information Criterion (AIC_c_; **Table 2**). In contrast, both simplified models provide better fits to female MLH1 counts than the more complex model with a distinct adaptive optimum for *M. m. musculus*. We recover qualitatively identical findings when modeling these evolutionary regimes along both NJ and ML trees (**Table 2**). For both trees, the maximum likelihood estimate of the optimum male trait value (𝜃_𝑚,𝑚𝑢𝑠𝑐_) for the *M. m. musculus* lineage is ∼3 MLH1 foci higher than that for background lineages, and estimates of the strength of selection toward the *M. m. musculus* adaptive optimum (𝛼) are at least an order of magnitude greater than that for the single optimum model. Taken together, these findings indicate that global recombination rates are evolving according to distinct evolutionary regimes in *M. musculus* males and females, and provide evidence for an adaptive, lineage-specific increase in male recombination rates in *M. m. musculus*.

**Table 2.** Evaluation of distinct phylogenetic models of sex-specific recombination rate evolution

| Tree | Sex | Model | $\alpha$ | $\sigma^2$ | $\theta_a$ | $\theta_{Mmm}$ | Likelihood | df | AIC - corrected |
| --- | --- | --- | --- | --- | --- | --- | --- | --- | --- |
| NJ | M | Brownian motion |  | 24.090 |  |  | -52.563 | 2 | 109.757 |
| NJ | M | Single optimum |  |  |  |  | -49.561 | 3 | 106.456 |
| NJ | M | *M. m. musculus* | 4486.79 | 38426.47 | 23.19 | 26.44 | -47.218 | 4 | 104.790 |
| NJ | F | Brownian motion |  | 8.34 |  |  | -24.041 | 2 | 53.416 |
| NJ | F | Single optimum | 8.99 | 44.66 | 25.30 |  | -22.424 | 3 | 53.847 |
| NJ | F | *M. m. musculus* | 9.98 | 49.32 | 25.23 | 25.55 |  | 4 | 58.508 |
| ML | M | Brownian motion |  | 35.54 |  |  | -51.561 | 2 | 107.754 |
| ML | M | Single optimum | 6.30 | 75.20 | 24.12 |  | -49.048 | 3 | 105.429 |
| ML | M | *M. m. musculus* | 1431.38 | 12265.74 | 23.19 | 26.44 | -47.218 | 4 | 104.790 |
| ML | F | Brownian motion |  | 19.35 |  |  | -26.35381 | 2 | 58.04095 |
| ML | F | Single optimum | 30.13 | 149.45 | 25.30 |  | -22.47142 | 3 | 53.94283 |
| ML | F | *M. m. musculus* | 322.28 | 1574.06 | 25.17 | 25.60 | -22.38619 | 4 | 58.48667 |

$\alpha$ : strength of directional force toward optimal trait value, units are inverse of time
$\sigma^2$ : magnitude of random perturbations tolerated about the optimal trait value, units measured as MLH1 foci x time
$\theta_a$ : ML estimate of optimal MLH1 count across the tree
$\theta_{Mmm}$ : ML estimate of optimal trait value along *M. m. musculus* lineage

To further confirm these findings, we next modeled the magnitude of sex dimorphism in MLH1 foci counts between females and males according to the same evolutionary regimes. Although our dataset is modest in size (n = 15 taxa), model fitting with the ML tree favors an evolutionary regime in which a distinct trait optimum is favored in *M. m. musculus* compared to other lineages (AIC_c_, musculus lineage = 68.34; AIC_c_, single optimum = 69.64). We obtain qualitatively identical results when modeling these evolutionary scenarios along the NJ tree (AIC_c_, musculus lineage = 68.34; AIC_c_, single optimum = 68.57; **Table S4**).

## DISCUSSION

The rate of recombination shapes outcomes of evolutionary processes in nature, but we lack a comprehensive understanding of how the molecular phenotype of recombination itself evolves (Johnston 2024). Prior work has established significant variation in recombination rate between species, within populations, and between males and females, but few studies have analyzed this phenotypic diversity in a phylogenetic framework. Here, we combined MLH1 foci counts across 15 genetically diverse inbred mouse strains with legacy datasets to evaluate the evolution of recombination rates across 31 wild-derived inbred house mouse strains in a sex-specific phylogenetic context. Our data confirm prior speculation that recombination rate evolution has proceeded under distinct evolutionary regimes in male and female house mice (Dumont and Payseur 2011; Peterson and Payseur 2021). We show that female recombination rates display no phylogenetic signal in *Mus*, are governed by a single phylogeny-wide adaptive optimum value, and exhibit a general trend of evolutionary stasis. In contrast, the phylogenetic distribution of male recombination rates is better approximated by the structure of the underlying phylogeny and has been influenced by a lineage-specific increase in *M. m. musculus*.

Curiously, elevated male recombination rates are observed in only a subset of our surveyed *M. m. musculus* strains (PWD/PhJ, PWK/PhJ, and Gor/TUA). Male recombination rates in other *M. m. musculus* strains (CZECHI/EiJ, CZECHII/EiJ, TOM/TUA, AST/TUA, and KAZ/TUA) are close, or even less than, the phylogeny-wide mean (**Figure 1**). Thus, the male-specific increase in recombination rate in *M. m. musculus* is not universal across this clade. These observations can be interpreted through the lens of mouse demographic history. Current models posit that the three core *M. musculus* subspecies emerged from a central population in southwestern Asia, and radiated westward, northward, and eastward very recently and nearly simultaneously (<0.5 MYA), giving rise to the *domesticus*, *musculus*, and *castaneus* subspecies lineages, respectively (Boursot et al. 1993b; Phifer-Rixey et al. 2020). As subspecies ranges expanded, they came into secondary contact, leaving substantial footprints of introgression and hybridization in contemporary wild mouse genomes (Fujiwara et al. 2022; Morgan et al. 2022). Further, because of their recent evolutionary origins, many loci in mouse genomes exhibit incomplete lineage sorting due to ancestral allele sharing between subspecies (White et al. 2009; Keane et al. 2011). Two of the high recombination rate *M. m. musculus* strains (PWD/PhJ and PWK/PhJ) derive from wild-caught mice sampled from close to a natural hybrid zone between *M. m. domesticus* and *M. m. musculus* that transects Europe (Macholán et al. 2012), while the wild-caught progenitors of the third *M. m. musculus* strain (Gor/TUA) derive from a more distant location in Siberia (Gorno-Altaisk, Russia). Future work will be required to determine whether the the extremely high recombination rate phenotype in these three geographically isolated strains (1) reflects the population sharing of alleles that emerged uniquely in *M. m. musculus* following subspecies migration out of the ancestral region, (2) is due to ancestral alleles that were subsequently lost in *M. m. castaneus* and *M. m. domesticus*, or (3) is attributable to recent introgression events between *M. m. musculus* populations, potentially aided by human migration and the status of house mice as human commensals.

Our findings raise a key, outstanding question: what are the biological mechanisms that drive the sex-specific evolution of recombination rate in house mice? On a genetic level, sex-specific recombination rates could be underlain by alleles with sex-specific, sex-limited, or sex-linked effects on recombination (Pennell and Morrow 2013; Mank 2017a; Mank 2017b). Indeed, prior linkage studies have identified numerous recombination rate modifiers, including alleles with opposite effects on recombination in males and females (Kong et al. 2008; Halldorsson et al. 2019), alleles with effects in only one sex (Kong et al. 2014; Ma et al. 2015; Brekke et al. 2023), and sex-linked recombination rate modifiers (Dumont and Payseur 2011; Liu et al. 2014). Our on-going efforts to define the genetic architecture of sex-specific recombination rates in *M. musculus* stand to provide further insight into the molecular underpinnings of this dimorphic and labile trait.

The discovery of a sex- and lineage-specific increase in recombination rate in *M. musculus* is especially exciting in the context of prior work on the genetic control of recombination in this system. Multiple large- and moderate-effect modifiers of male recombination rate have been mapped to the X chromosome and autosomes in intersubspecific crosses between *M. m. musculus* and *M. m. castaneus*, with X-linked and autosomal modifiers exhibiting antagonistic effects (Dumont and Payseur 2011; Liu et al. 2014; Dumont 2017b). Specifically, in males, X-linked alleles from the low recombination rate subspecies (*M. m. castaneus*) confer an increase in recombination rate, whereas autosomal alleles from *M. m. castaneus* are associated with recombination rate decreasing effects. One of these X-linked modifiers overlaps a locus implicated in hybrid male sterility in intersubspecific crosses between *M. m. musculus* and *M. m. domesticus*, suggesting a possible link between recombination and the emergence of nascent subspecies barriers (Balcova et al. 2016). More recent work has demonstrated that a genetic modifier of female recombination rates is also resident on the mouse X chromosome (Liu et al. 2014; Balcova et al. 2016; Dumont 2017b), although additional work is required to demonstrate that the loci identified in males and females colocalize.

From an evolutionary perspective, support for unique sex-specific evolutionary programs implies that natural selection favors distinct recombination rate optima in males and females. Although aberrant recombination profiles are frequently associated with aneuploidy or infertility (Hassold and Hunt 2001; Oliver et al. 2008; Lu et al. 2012; Bell et al. 2020; Carioscia et al. 2025), the relationship between natural variation for recombination rate and reproductive fitness remains poorly understood (Ritz et al. 2017). Quantitative genetic studies have established that natural variation for recombination rate is shaped by adaptation in wild *Drosophila* populations (Samuk et al. 2020), but further direct evidence remains scant. Human females with higher recombination rates birth more children (Kong et al. 2004; Campbell et al. 2015), but it is not clear whether this correlation reflects the increased fitness of females with higher recombination rates or the increased odds a fertilized oocyte with elevated recombination rate survives early development in older mothers. Further, to our knowledge, there exists no evidence that male recombination rates are linked to individual differences in fitness.

Unfortunately, evaluating the possibility that the sex-specific variation in MLH1 foci counts observed across *M. musculus* is linked to fitness is not currently possible as we lack estimates for fitness-associated traits for all but a few of the strains surveyed in our study. Colony breeding records document strain-specific breeding performance and can provide a read-out of strain fitness. However, the reproductive performance of inbred strains is an amalgamation of male and female factors that cannot be readily disentangled to permit sex-specific estimates of fitness-associated traits. Further, the interpretation of fitness in inbred strains is likely confounded by the inbreeding process itself, which is associated with the fixation of deleterious variants at a rate determined by the mutation load of the parent population. These complexities call attention to the need for studies that profile sex-specific recombination rates and fitness measures in natural or outbred populations.

Although we lack direct experimental evidence for sex differences in recombination rate fitness optima, our finding that the evolution of recombination in house mice has unfolded according to sex-specific evolutionary regimes is strongly consistent with this possibility. Sex differences in trait fitness optima impose an inherent genetic conflict, as an allele that advances the trait value in the direction favoring one sex will have a negative effect on the fitness of the other sex. Such conflict can be genetically resolved via intralocus mechanisms, including sex-limited gene expression (Mank 2017a), sex-biased epigenetic silencing (Pennell and Morrow 2013), dominance reversal (Barson et al. 2015), gene duplication (VanKuren and Long 2018), or sequestration of genes with sexually antagonistic fitness effects to the sex chromosomes (Roberts et al. 2009; Mullon et al. 2012). Alternatively, sexual conflicts can be resolved via interlocus mechanisms involving distinct loci with opposing fitness effects in the two sexes. This latter scenario can fuel an evolutionary arms race between the sexes, wherein male-beneficial (female-detrimental) alleles impose a selective advantage for the emergence of female-beneficial (male-detrimental) alleles elsewhere in the genome (Schenkel et al. 2018). Overall, genetic conflict can maintain exceptionally high levels of trait polymorphism, propel rapid trait divergence, and shape key aspects of trait architecture (Mank 2017b; Schenkel et al. 2018).

These telltale hallmarks of interlocus genetic conflict are evident in the (1) high levels of recombination rate polymorphism within house mouse subspecies, (2) unique, sex-specific patterns of accelerated recombination rate divergence across *Mus*, and (3) the genetic control of recombination by X-linked and autosomal genetic loci with antagonistic effects in *M. musculus*. Thus, our findings layer new evidence onto the hypothesis that recombination rate variation across house mice may be driven, at least in part, by genetic conflict (Dumont 2017b). Additional work is clearly required to establish the validity of this hypothesis and identify the aspects of meiosis that may be subject to conflicting evolutionary pressures in males and females. Further, whether genetic conflict is a common mechanism shaping the evolution of recombination rates across taxa, or a potential phenomenon specific to house mice, warrants future experimental testing.

## DATA AVAILABILTY

The MLH1 foci count data analyzed in this report are provided in **Table S1**. File S1 contains the phylogenetic trees used for phylogenetic analysis and model fitting. Raw fastq sequencing data used in phylogenic reconstruction are available at the accession numbers listed in **Table S3**. Whole genome sequence data for the three strains newly sequenced as part of this study POHN/DehMmJax, Gor/TUA, and GAIB/NachJ are provided on the SRA Archive under PRJNA1276540. All supplemental figures, tables, and R code used to perform statistical and phylogenetic analyses can be found at https://figshare.com/projects/Recombination_Rates_Have_Been_Shaped_by_Sex-Specific_Evolutionary_Programs_in_House_Mice/252902.

## ACKNOWLEDGEMENTS

We gratefully acknowledge the contribution of the Genome Technologies Scientific Services at The Jackson Laboratory for their contributions to this work. We also thank members of the Research IT team at The Jackson Laboratory for providing access and support to the high-performance computing resources needed to perform this study. Lastly, we thank members of the Dumont Lab for helpful discussions and feedback on this project. This work was supported by funds from an NSF CAREER Award (DEB1942620) to BLD.

